# A conserved peptidase governs glucose homeostasis in *Bacteroides*

**DOI:** 10.64898/2026.04.10.717545

**Authors:** Kamalesh Verma, Seth G. Kabonick, Kailyn M. Winokur, John M. Flanagan, Eduardo A. Groisman, Guy E. Townsend

## Abstract

Glucose homeostasis is governed by peptidases across the kingdom of life. Host overconsumption of glucose transforms gut microbial compositions and activities. Here, we identify an IDE-like M16 peptidase in human gut commensal *Bacteroides* that controls glucose-dependent inhibition of the transcriptional regulator, Cur, which is important for intestinal colonization and host-microbial interactions. This regulatory paradigm is independent of fructose, establishing a specific pathway whereby glucose signaling hinders commensal fitness in the host. We determined that this peptidase cleaves targets of other M16-family peptidases, including insulin, to control carbon metabolism through the oxidative pentose phosphate pathway (OPPP). Furthermore, we show that peptidase activity governs the abundance of glycolytic enzymes to alter Cur activity. These findings establish that the activity of an M16 peptidase mediates changes to global transcription in *Bacteroides* species when glucose is abundant in the host diet.

## Introduction

Peptidases regulate glucose homeostasis in mammals and their resident microbiota^1–3^. Humans systemically control blood glucose levels using the antagonistic peptide hormones insulin and glucagon, which work in opposition to balance glucose uptake during feasting and mobilization during fasting^4,5^. The abundance and activity of both peptides are governed by sequential proteolytic events that first convert pro-hormone precursors into active forms, then degrade hormonal signals using the M16 class peptidase insulin-degrading enzyme (IDE)^1,2^. Insulin binding to cell-surface receptors prompts intracellular metabolic and post-translational cascades to alter transcription factor activity^6^. Many gut commensal bacteria from the *Pseudomonadota* or *Bacillota* phylum also possess and employ M-class peptidases to regulate the expression of glucose-dependent phospho-transferase systems (PTS), which phosphorylate glucose to initiate glycolysis and alter intracellular transcription^3^. The dominant intestinal phylum *Bacteroidota* do not possess PTS orthologs and instead use ATP- and pyrophosphate (PPi)-dependent glycolytic pathways to couple glucose catabolism with transcriptional responses^7,8^. In this work, we identify an M16 peptidase in *Bacteroides* that controls the abundance of glycolytic enzymes during glucose fermentation in the human gut, indicating that mammals and commensal bacteria rely on a conserved peptidase family for glucose homeostasis.

Prolonged abundant sugar consumption impacts human health by promoting insulin resistance, non-alcoholic fatty liver disease, and metabolic syndrome in the host^9,10^. Host overconsumption of glucose can also disrupt the fitness and alter the activities of commensal bacteria ^11,12^. Excessive dietary monosaccharides can overwhelm intestinal transporters and escape into the distal gut, exposing microbes to carbohydrates that were traditionally encountered as polysaccharides^13^. *Bacteroides* species grow logarithmically on monosaccharides *in vitro*, but their abundance is reduced *in vivo* when the host consumes glucose-rich diets^14,15^. Subsequent glucose fermentation by resident microbes is sufficient to drive intestinal colitis in mice^16,17^, establishing that sugar-induced changes in bacterial products can promote disease independent of host metabolic dysfunction.

Glucose and fructose reshape the transcriptome of the commensal gut bacterium *Bacteroides thetaiotaomicron* (*Bt*) by inhibiting the master regulator of carbohydrate utilization Cur, which is necessary for metabolizing a wide array of host- and diet-derived carbohydrates, including fucose, xylose, and ribose in the lumen and mucosa^18^. Cur is a DNA-binding transcriptional regulator that controls 20% of the *Bt* transcriptome, extending beyond carbohydrate utilization to include genes important for nutrient availability, host interactions, and intestinal colonization^19^. For example, Cur indirectly regulates the abundance of Roc, a critical fitness determinant *in vivo*^20,21^, and EF-G2, which facilitates GTP-independent protein synthesis to conserve energy during colonization^19,22^. Cur also coordinates the presentation of BT4295, a T-cell antigen that decorates outer membrane vesicles to promote differentiation^17^. In *Bacteroides* species, the catabolism of glucose and fructose through ATP-dependent glycolysis reduces intestinal fitness by silencing *cur* - dependent gene products^8^. Consequently, genetic disruption of this pathway partially restored Cur activity when simple-sugars were present, but does not completely alleviate inhibition, suggesting that additional regulators govern Cur in *Bt*.

Here, we developed a forward genetic screen that identified an IDE-like, M16 peptidase encoded by *BT3803* that regulates Cur activity during glucose catabolism. We show that this peptidase forms a catalytic heterodimer with the distally encoded gene product that can proteolytically act on established peptide substrates of M16 orthologs across kingdoms, including insulin. We found that peptidase activity is dispensable for Cur inhibition by fructose, unlike the ATP-dependent phosphofructokinase^8^, establishing distinct regulatory pathways for these isomeric dietary sugars despite similar catabolic routes. RNA-seq reveals that proteolytic activity alters ∼700 *cur* -dependent and -independent transcripts in *Bt*. Remarkably, a mutant strain that lacks this peptidase exhibits a competitive advantage over *wild-type Bt* during *in vitro* growth in glucose and in gnotobiotic mice provided a concentrated glucose beverage, indicating that host consumption of simple sugars disrupts commensal fitness. Using proximity labeling, we identified numerous proteinprotein interactions, including many that mediate glycolysis and nucleotide metabolism. We show that the proteolysis of key metabolic enzymes, including the ATP-producing glycolytic enzyme pyruvate kinase, is important to regulate their abundances and control Cur activity. Collectively, these findings uncover a regulatory paradigm that connects M16 peptidase activity to global transcription during *Bacteroides* glucose catabolism in the mammalian gut.

## Results

### Identification of genes that inhibit Cur activity in *Bacteroides*

To identify factors that facilitate glucose-dependent Cur inhibition, we generated a transposon insertion library in a *Bt* strain expressing an epitope-tagged Roc protein. We leveraged Roc as readout of Cur activity because it was successfully used to identify *trans*- and *cis*-regulatory elements mediating *cur* - dependent responses to nutrient availability ^20,21^. We examined 5,000 colonies on solid minimal media containing glucose by colony blotting and isolated 24 strains that produced Roc amounts greater than *wild-type*. Sequencing revealed that 22 mutants harbored unique insertion sites within the *Bt* genome that corresponded to three distinct loci (Figure 1A): *BT2062*, encoding an ATP-dependent phosphofructokinase; *BT3802-3*, encoding a putative *cis*/*trans*-prolylisomerase and an M16-family peptidase, respectively; and *BT0173-4*, encoding a GGGtGRT protein and a NifU homolog, respectively.

**Figure 1.**
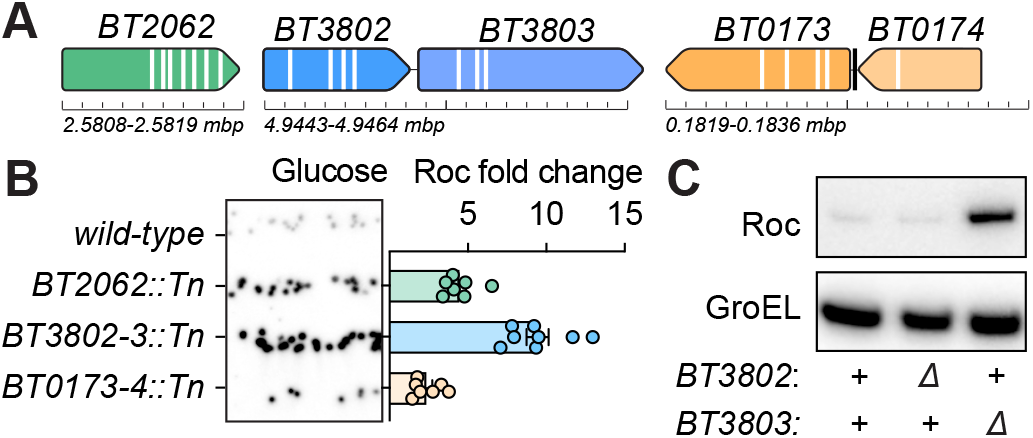
Identification of mutants defective for Cur inhibition in glucose. **A** Schematic depicting genomic location and transposon insertion sites in the three loci identified using a forward genetic screen. **B** Colony blot of Roc protein amounts from *wild-type, BT2062*::Tn, *BT3802-3*::Tn, or *BT0173-4*::Tn on solid media containing glucose (left); Fold change of Roc protein amounts relative to *wild-type* from transposon mutants identified in the genetic screen grown in liquid media containing glucose (right). **C** Western blot of Roc protein amounts from *wild-type*, Δ*BT3802* or Δ*BT3803* grown in minimal media containing glucose. For **B**, points represent three independent measurements of individually isolated transposon mutants.

Transposon mutants were further characterized by measuring bacterial growth and quantifying Roc protein amounts in liquid media containing glucose. Surprisingly, strains harboring transposon insertions in each of the three genetic loci did not exhibit reduced growth compared to *wild-type*, suggesting that these genes are metabolically dispensable *in vitro* (Supplementary Figure 1A). Strains harboring transposons corresponding to *BT2062* (*pfkA*) exhibited 4.5-fold increased Roc, consistent with our previous findings that ATP-dependent phosphofructokinases, such as PfkA, are required for glucose-dependent reduction in Cur activity and validating our genetic screen (Figure 1B)^8^. Mutants with insertions in the *BT3802-3* locus produced 10-fold increased Roc protein levels, whereas *BT01734* insertions resulted in a modest 2-fold increase (Figure 1B). Here, we focused our investigation on *BT3802-3* because these mutants produced the greatest fold increases in Roc amounts and we previously established the role of PfkA in controlling Cur when glucose or fructose is available ^8^.

A single putative transcription start site precedes the *BT3802-3* locus^23^, indicating that these genes are cotranscribed. We constructed strains lacking ORFs of either gene to understand how each contributes to glucose-dependent Roc silencing. Accordingly, insertions in *BT3802* are polar on downstream transcription because only a *BT3803*-deficient strain (Δ*mdpA*) exhibited increased Roc amounts compared to *wild-type* (Figure 1C & Supplementary Figure 1B). Introduction of a complementing plasmid into a *BT3803* mutant restored Roc protein levels to those resembling the *wild-type* strain (Supplementary Figure 1B). Thus, *Bt* requires a predicted metal-dependent peptidase (*mdpA*) to silence Roc production during growth in glucose.

### MdpA forms a heterodimeric M16 peptidase complex

A phylogenetic analysis of M16 peptidases revealed that MdpA is conserved across genera throughout *Bacteroidota*, including *Prevotella, Porphyromonas*, and *Bacteroides*, suggesting that this peptidase is an ancestral feature rather than a recent acquisition in *Bt* (Figure 2A). Sequence alignment of MdpA with other peptidases, such as Sph2681 from *Sphingomonas* A1^24^, BH2405 from *Halalkalibacterium halodurans*^25^, and PtrA from *Escherichia coli* (*Ec*)^26^ revealed that MdpA shares a conserved HXXEH motif required for metal cofactor binding and proteolytic activity (Figure 2B & Supplementary Figure 1C). This motif is characteristic of M16 family peptidases, which are comprised of distinct structural classes: monomers with two fused domains (e.g., PtrA and IDE)^2^, homodimers (e.g., BH2405)^25^, or heterodimers (e.g., Sph2681)^24^.

**Figure 2.**
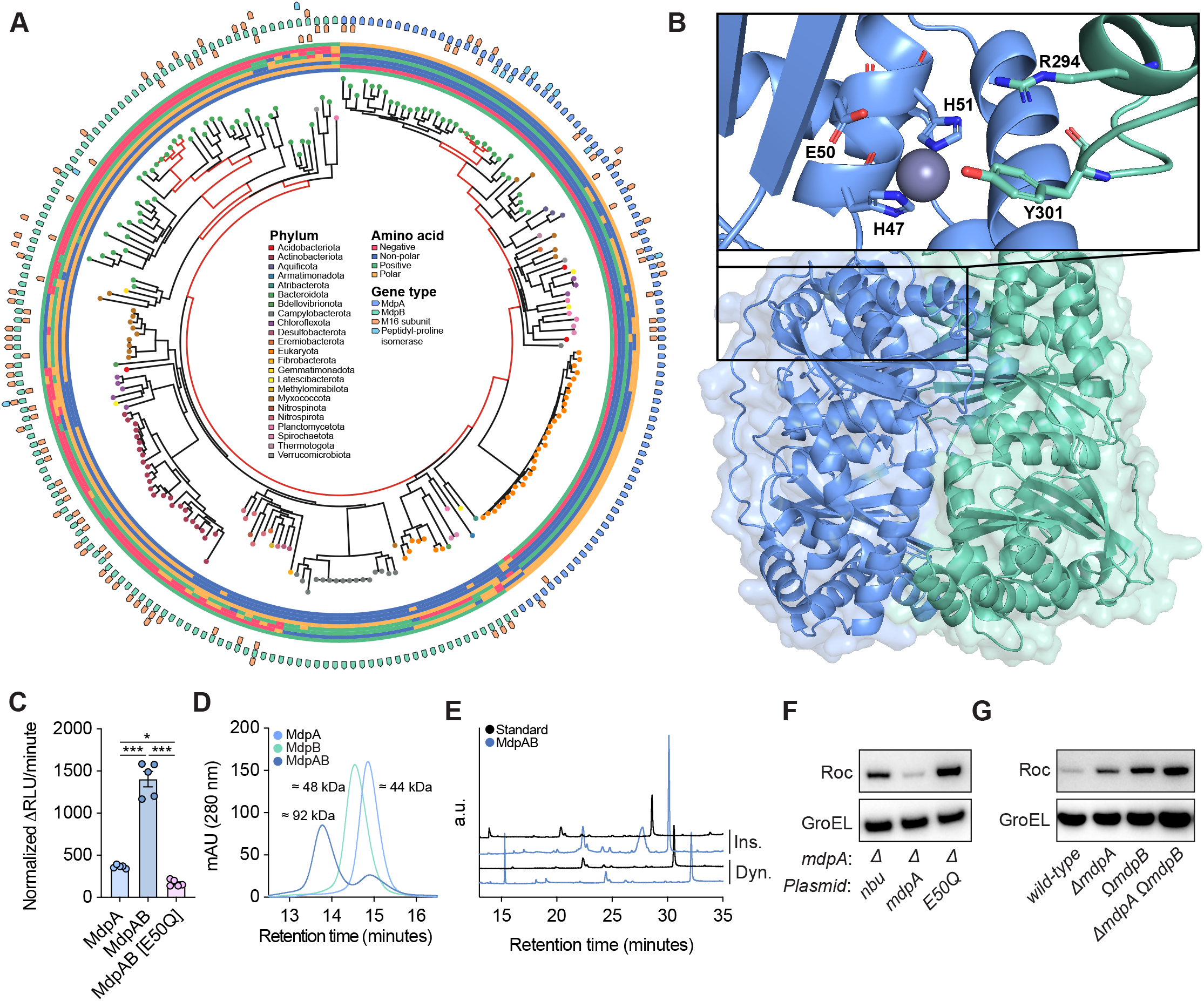
MdpA belongs to a widely conserved M16 peptidase family. **A** Phylogenetic analysis of the distribution of M16 class peptidases across prokaryotic and eukaryotic species. Branches corresponding to the *Bacteroides* genus are colored in red. **B** Structural alignment of an AlphaFold-predicted MdpA (blue) and MdpB (green) complexed with a zinc ion. **C** Peptidase assay measuring the proteolytic cleavage of the fluorogenic substrate, Abz-pAK. **D** Size-exclusion chromatogram of MdpA, MdpB, or equal amount of MdpAB. **E** Liquid chromatography analysis of insulin chain B or dynorphin A peptide cleavage with or without MdpAB. **F** Western blot of Roc amounts from *wild-type* or Δ*mdpA* harboring an empty vector (nbu) or plasmid-borne copies of *mdpA* or E50Q grown in minimal media containing glucose. **G** Western blot of Roc amounts from *wild-type*, Δ*mdpA*, Δ*mdpB*, and Δ*mdpA* Δ*mdpB* grown in minimal media containing glucose. For **C**, N=8 technical replicates; error represent SEM. P-values were calculated using one-way ANOVA and * represents values *<* 0.001.

MdpA lacks fused domains characteristic of the architectures typically found in monomeric M16 peptidases. Moreover, purified MdpA exhibited poor cleavage of a fluorophore-labeled peptide derived from the N-terminal region of adenylate kinase (Abz-pAK), a known substrate of M16-family members (Figure 2C)^24^, implying that MdpA exists as a heterodimer and requires an additional subunit for activity. Although partner subunits of M16 peptidases are frequently encoded in the same operon (Figure 2A)^24,27^, the genomic context of *mdpA* revealed no reasonable candidates. Instead, *Bacteroidota* genera encode *mdpA* alongside a putative *cis*/*trans*-prolylisomerase (e.g., *BT3802* in *Bt*) that is not involved in Roc silencing (Figure 1C & Figure 2A). To identify a candidate heterodimer partner, we computationally screened *Bt* for orthologs of Sph2682, the non-catalytic subunit that complexes with Sph2681 for proteolytic activity in *Sphingomonas* A1^24^. BT0746 (MdpB) emerged as a strong candidate because it contains a hallmark R(X)_3,5_Y domain conserved in M16 heterodimers and forms a high-confidence AlphaFold3 prediction when modelled with MdpA (ipTM of 0.92; pTM of 0.93) (Figure 2A, B)^28^. Dimer formation was tested by recombinantly expressing both subunits individually in *Ec* and subjecting each to size exclusion chromatography. When tested alone, MdpA and MdpB eluted as monomers with molecular weights of 44 kDa and 48 kDa, respectively. When combined in equal amounts, the two subunits preferentially formed a 92-kDa complex (MdpAB) (Figure 2D). Additionally, MdpAB exhibited strong catalytic activity against established non-native substrates, including Abz-pAK, insulin chain B, and dynorphin A (Figure 2C, E).

To determine whether the catalytic activity of MdpA is required for glucose-dependent Roc silencing, we expressed an MdpA allele harboring an E50Q point mutation within the conserved HXXEH motif. The E50Q mutant complexed with MdpB lacked strong *in vitro* activity against Abz-pAK (Figure 2C), exhibiting half the activity of MdpA alone. Consistent with this, glucose failed to decrease Roc amounts in cells expressing the E50Q mutant, whereas complementation with *wild-type mdpA* restored Roc silencing and *cur*-dependent transcripts to levels resembling the parental strain (Figure 2F & Supplementary Figure 1B, D, E). Because both peptidase subunits are required for activity, we reasoned that *mdpB* is also necessary for glucose-mediated Roc silencing. Indeed, disabling *mdpB* (Ω*mdpB*) phenocopied Roc amounts produced by Δ*mdpA* (Figure 2G), demonstrating that the complete MdpAB heterodimer complex is necessary for both peptidase activity and Cur inhibition during glucose catabolism.

### MdpA specifically facilitates Cur inhibition by glucose

Roc production is controlled directly through *cis*-acting mRNA leader residues and indirectly by Cur^20^. We reasoned that if MdpA decreases Roc abundance by inhibiting Cur activity, *mdpA* inactivation should increase expression of *cur*-dependent genes. An *mdpA*-deficient strain exhibited at least 2-fold changes in 543 previously established *cur*-dependent transcripts^19^, including 2-fold and 3-fold increased expression of *fusA2* and the T-cell antigen *BT4295*, respectively (Figure 3A), revealing that loss of MdpA broadly alters transcription by controlling Cur activity in this condition. We also detected 139 transcripts outside the known Cur regulon, demonstrating that MdpA also controls additional *cur*-independent genes. The latter included the *BT3571-5* locus that exhibited strong, statistically significant upregulation in Δ*mdpA*. However, a *BT3571-5* deficient strain did not exhibit changes in Roc or Cur regulated transcripts (Supplementary Figure 3A, B, C), indicating that this locus does not control Cur activity. A double-mutant strain lacking both *mdpA* and *cur* (Δ*mdpA* Δ*cur*) exhibited Roc amounts that were indistinguishable from a *cur* mutant (Δ*cur*) in glucose (Figure 3B & Supplementary Figure 3A). Similarly, the *cur*-dependent transcripts *fusA2* and *BT4295* were reduced when these strains were grown in glucose (Figure 3C & Supplementary Figure 3B, respectively). Finally, complementation with plasmid-encoded *cur* restored Roc and *cur*-dependent transcript levels in Δ*cur*, but complementation with *mdpA* did not (Figure 3B, C), establishing that MdpA inhibits Cur to silence Roc production during growth in glucose.

**Figure 3.**
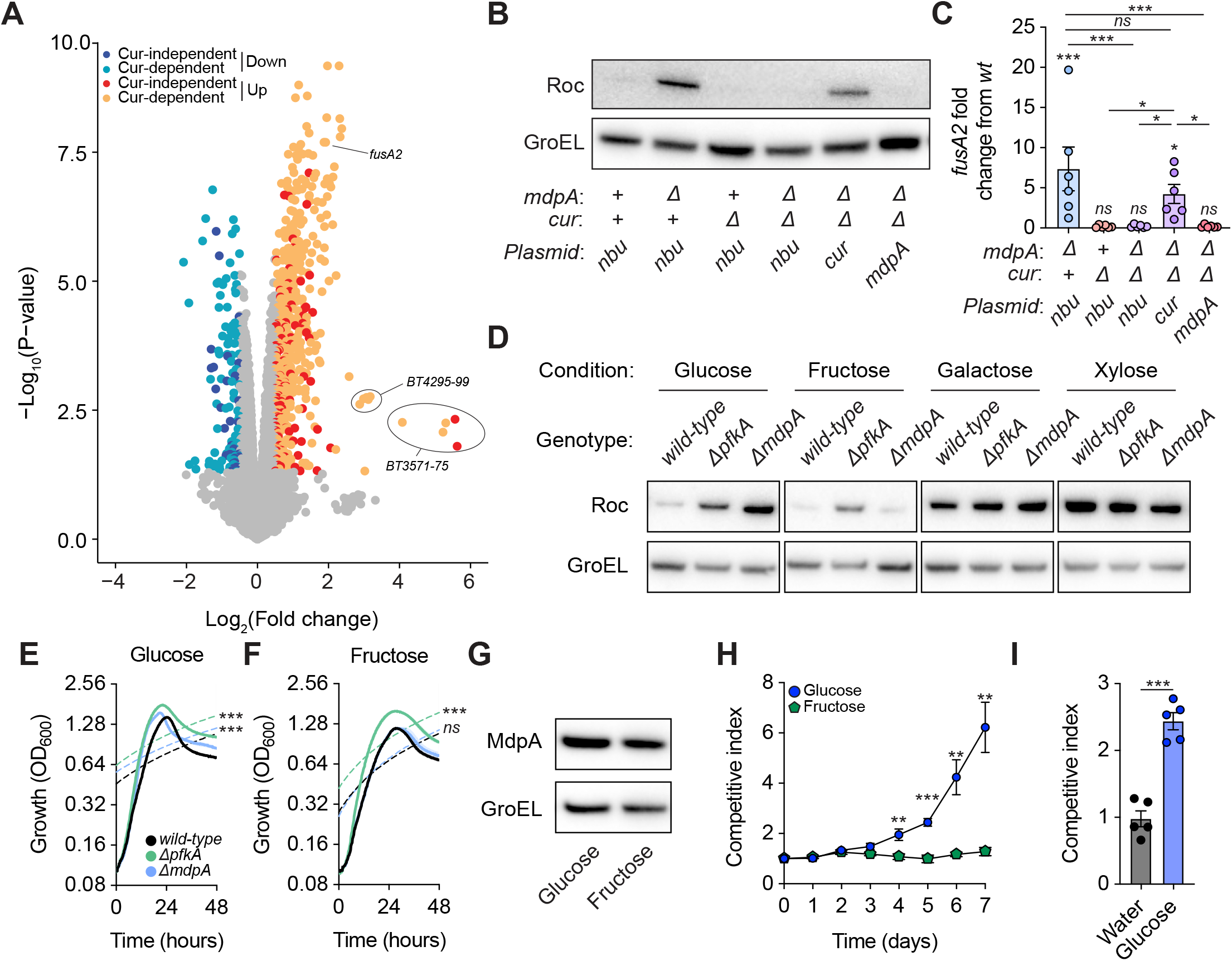
MdpA is required for Cur inhibition in glucose but not fructose. **A** Volcano plot depicting differentially abundant transcripts in Δ*mdpA* compared to *wild-type* in minimal media containing glucose. **B** Western blot of Roc amounts from *wild-type*, Δ*mdpA*, Δ*cur*, or Δ*mdpA* Δ*cur* harboring an empty vector (nbu), *mdpA* or *cur* grown in minimal media containing glucose. **C** Fold change of *fusA2* transcript amounts relative to *wild-type* from Δ*mdpA*, Δ*cur*, or both harboring an empty vector (nbu), *mdpA*, or *cur* grown in minimal media containing glucose. **D** Western blot of Roc amounts from *wild-type*, Δ*pfkA*, or Δ*mdpA* mutants grown in minimal media containing glucose, fructose, galactose, or xylose. **E, F** Growth of *wild-type*, Δ*pfkA*, or Δ*mdpA* mutants grown in minimal media containing glucose (**E**) or fructose (**F**). **G** Western blot of MdpA protein amounts from *wild-type Bt* in minimal media containing glucose or fructose. **H** Competitive index of Δ*mdpA* compared to *wild-type* in glucose or fructose containing rich media. **I** Fitness of Δ*mdpA* compared to *wild-type* in gnotobiotic mice provided water or a 30% glucose solution. For **C**, N=5 biological replicates; error bars represent SEM. For **E, F**, N=8 biological replicates; error represents SEM in color matched shading. For **H, I**, N=5 biological replicates; error bars represent SEM. For **C, E**, & **F** P-values were calculated using one-way ANOVA and * represents values *<* 0.05, ** *<* 0.01, *** *<* 0.001. For **H**, P-values were calculated using two-way ANOVA and ** represents values *<* 0.01, *** *<* 0.001. For **I**, P-values were calculated using an unpaired t-test and *** represents values *<* 0.001.

We examined whether MdpA inhibited Cur activity during growth in glucose or fructose, similar to PfkA^8^. Unexpectedly, Δ*mdpA* produced Roc amounts comparable to *wild-type Bt* when grown in fructose, galactose, or xylose, whereas Δ*pfkA* exhibited 4-fold increased Roc amounts in both glucose and fructose (Figure 3D & Supplementary Figure 3C). Additionally, Δ*mdpA* exhibited an increased growth rate and maximum only in glucose compared to *wild-type Bt*, whereas Δ*pfkA* growth increased in both glucose and fructose (Figure 3E, F)^8^. This condition-dependent regulation is likely governed metabolically, rather than altering MdpA amounts in the cytoplasm, because HA-epitope tagged protein amounts were unaltered between carbohydrate sources (Figure 3G and Supplementary Figure 2D). Thus, each monosaccharide inhibits Cur activity through distinct mechanisms despite their isomeric structures, adjacent metabolic entry points, and similar catabolic pathways.

Because MdpA specifically inhibits *cur*-dependent genes important for bacterial fitness and carbohydrate utilization when glucose is present, we predicted that Δ*mdpA* would exhibit a fitness advantage in glucose but not fructose. Accordingly,Δ*mdpA* outcompeted *wild-type* when both strains were cocultured in glucose-containing media and serially passaged for 7 days, but the relative abundance of Δ*mdpA* and the *wild-type* strain remained similar in media containing fructose (Figure 3H). To determine whether Δ*mdpA* exhibited a similar advantage over *wild-type in vivo*, we introduced equivalent amounts of each strain into gnotobiotic mice supplied either normal drinking water or a 30% glucose solution. Remarkably, mice consuming the glucose-rich beverage harbored 2.5-fold more Δ*mdpA* relative to *wild-type* after 14 days, whereas mice provided water contained indistinguishable ratios between strains (Figure 3I). Taken together, these data demonstrate that MdpA is a primary regulator of Cur and that its peptidase activity reduces *Bt* intestinal fitness during host glucose consumption.

### MdpA-mediated inhibition of Cur requires the oxidative pentose phosphate pathway

In addition to degrading their substrates, M16 peptidases can post-translationally modify target proteins^25^; however, MdpA does not modify Cur during growth in glucose because Δ*mdpA* exhibited Cur protein amounts and migration indistinguishable from that in *wild-type Bt* (Supplementary Figure 4A). Because Δ*mdpA* exhibited *cur*-dependent effects only in glucose, we predicted that MdpA could alternatively target enzymes specific to glucose metabolism. The glucokinase (Glk, BT2493) in *Bt* is necessary for glucose-mediated Cur inhibition ^8^ but there were no differences in Glk amounts or migration in a Δ*mdpA* strain (Supplementary Figure 4B). Likewise, there were no differences observed for enzymes catalyzing three other major metabolic branch points downstream of glucose-6-phosphate (G6P): BT1548 (Pgm), which converts G6P to glucose-1-phosphate (G1P) for glycogenesis, BT2124 (Pgi), which converts G6P to fructose-6-phosphate (F6P) for glycolysis, and BT1221 (G6PD), which converts G6P to 6-phosphogluconolactone (GL6P) in the oxidative pentose phosphate pathway (OPPP) (Supplementary Figure 4B).

Because MdpA does not target these enzymes directly, we hypothesized that a downstream component of glucose catabolism was responsible for Cur inhibition (Figure 4A). Despite successfully engineering chromosomal deletions of *pgi* and *pgm* using fructose-rich media, these mutants exhibited severe growth defects in minimal media containing glucose, precluding genetic examination of their role in controlling Cur activity (Supplementary Figure 4C). Thus, we focused on three enzymes in the OPPP because it is required for Cur activation following carbon starvation and mutants lacking these enzymes grew similarly to the wild-type strain on glucose, rendering the OPPP genetically tractable ^20^. A strain lacking both *mdpA* and *pgl* (Δ*mdpA* Δ*pgl*) displayed 2.2-fold reduced Roc levels compared to Δ*mdpA* alone (Figure 4B & Supplementary Figure 4D). The *cur*-dependent transcripts *fusA2* and *BT4295* (Figure 4C and Supplementary Figure 4E) also exhibited similar trends, demonstrating that the OPPP is required for increased Cur activity in an *mdpA*-deficient strain.

**Figure 4.**
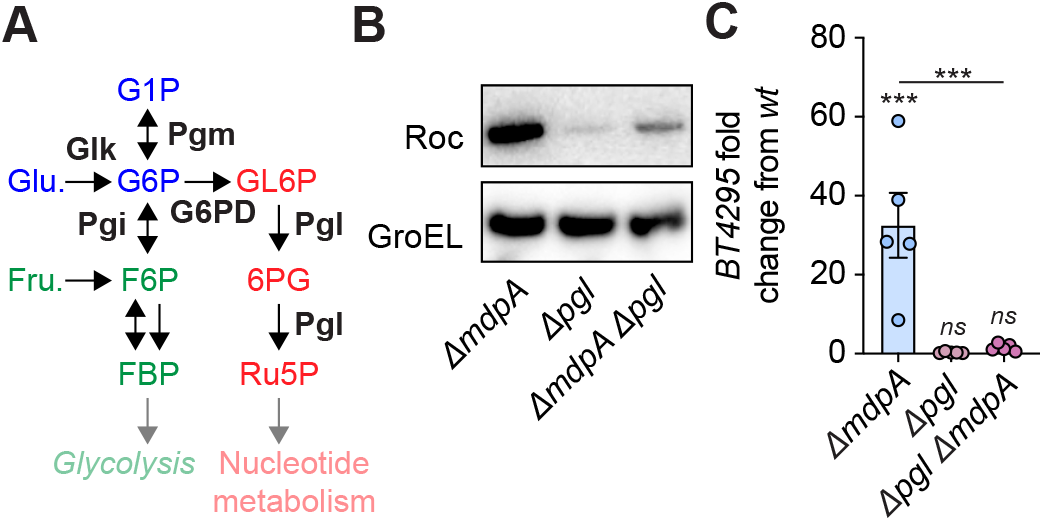
The oxidative pentose phosphate pathway is required for MdpA control of Cur activity. **A** Schematic of the glycolytic and oxidative pentose phosphate (OPPP) pathways in *Bacteroides*. **B** Western blot of Roc amounts from Δ*mdpA*, Δ*pgl*, or Δ*pgl* Δ*mdpA* grown in minimal media containing glucose. **C** Fold change of *BT4295* transcript amounts relative to *wild-type* from Δ*mdpA*, Δ*pgl*, and Δ*mdpA* Δ*pgl* grown in minimal media containing glucose. For **C**, N=5 biological replicates; error bars represent SEM. P-values were calculated using one-way ANOVA and * represents values *<* 0.05, ** *<* 0.01, *** *<* 0.001.

### Identification of MdpA interacting partners

To identify candidate proteolytic targets of MdpA, we developed a proximity labeling approach using BioID^29^, which leverages the activity of *Ec* BirA to biotinylate lysine residues on interacting partners. We constructed anhydrotetracycline (aTc)-inducible plasmids harboring *wild-type mdpA* or the catalytically inactive E50Q mutant fused to BirA (Figure 5A). The fusion protein retained the ability to inhibit Cur activity because Roc amounts were indistinguishable between Δ*mdpA* strains expressing either MdpA-BirA or native MdpA in the presence of aTc (Figure 5B). Furthermore, aTc addition was required to detect either fusion protein (Figure 5B) and without aTc, Roc levels in Δ*mdpA* resembled a strain harboring the empty vector (Supplementary Figure 5A, B). In contrast, strains expressing either the E50Q mutant or the E50Q-BirA fusion protein produced Roc resembling Δ*mdpA* harboring an empty vector, regardless of aTc addition (Figure 5B & Supplementary Figure 5A). These data establish that the MdpA-BirA fusion protein retains the catalytic activity necessary to inhibit Cur during growth in glucose and that BirA does not interfere with MdpA function.

**Figure 5.**
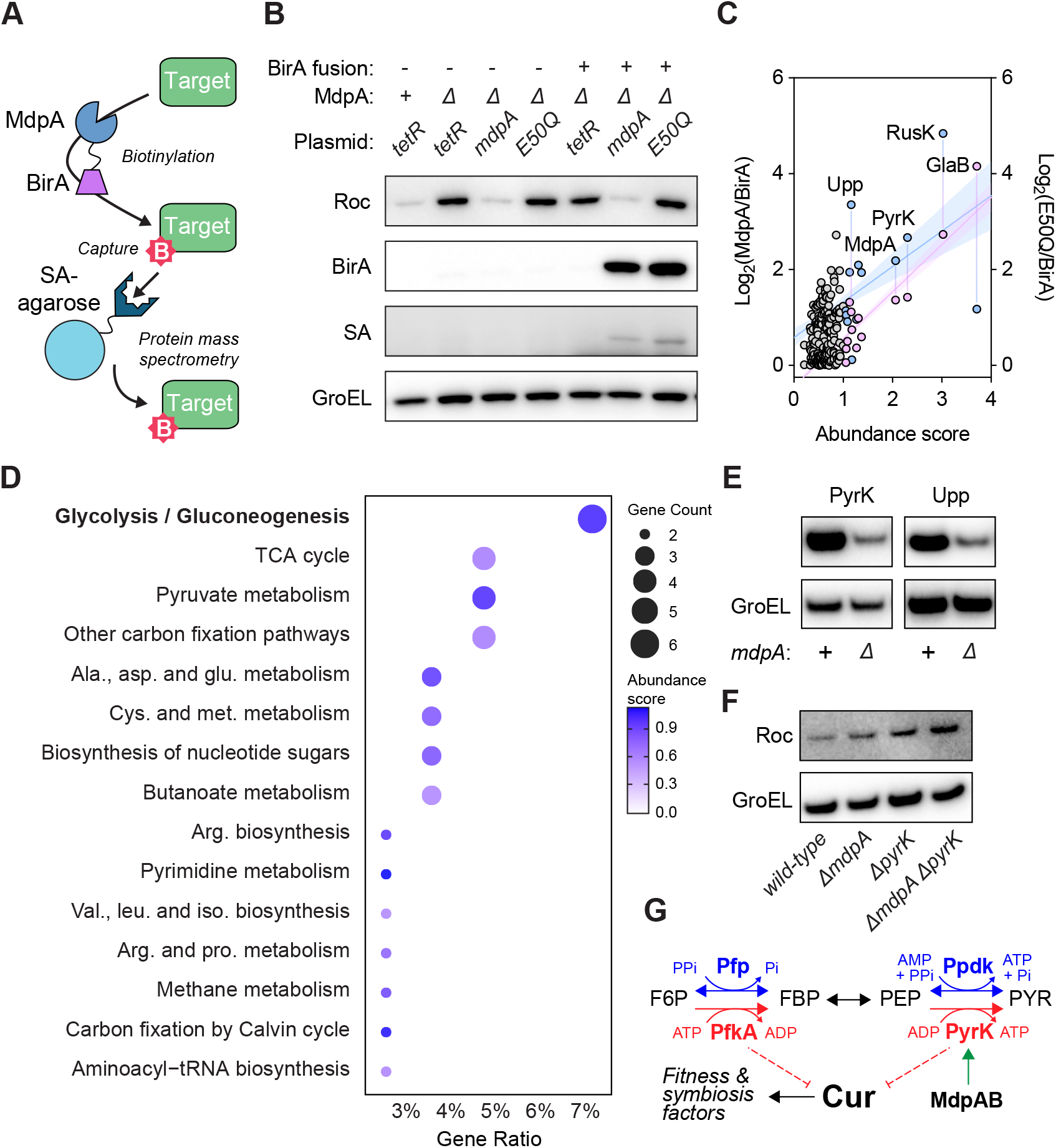
Identification of MdpA targets. **A** Schematic of BirA-dependent biotinylation of MdpA targets (SA: streptavidin). **B** Western blot of Roc, BirA, and SA protein amounts following aTc-inducible expression of MdpA or E50Q-BirA fusion proteins from *wild-type* or Δ*mdpA* grown in minimal media containing glucose. **C** Proteomic analysis of captured proteins reflected by Log_2_ fold change of MdpA or E50Q fusion proteins normalized by BirA. **D** KEGG pathway enrichment analysis of proteins identified following mass spectrometry of BioID proximity ligation and purification. **E** Western blot of PyrK (left) or Upp (right) amounts from *wild-type* or Δ*mdpA* grown in minimal media containing glucose. **F** Western blot of Roc amounts from *wild-type*, Δ*mdpA*, Δ*pyrK*, or Δ*mdpA* Δ*pyrK* grown in minimal media containing glucose. **G** Model schematic depicting glycolytic control of Cur activity in *Bacteroides*.

We performed streptavidin affinity purification from strains expressing MdpA-BirA, E50Q-BirA, or BirA alone (Figure 5A) and used mass spectrometry to identify biotinylated proteins. As expected, MdpA was highly enriched in the proteomic analysis for experiments using either fusion construct, indicating that BirA readily labeled its partner protein (Figure 5C). A gene ontology (GO) analysis of captured proteins with at least 10% coverage from strains expressing MdpA-BirA or E50Q-BirA fusion proteins compared to BirA alone identified many enzymes involved in glycolysis and nucleotide metabolism (Figure 5D). Among these was BT2841 (PyrK), the sole putative pyruvate kinase in *Bt*, which catalyzes the conversion of ADP and PEP to ATP and pyruvate (PYR). Intriguingly, PyrK is three reversible, metabolic steps downstream from the ATP-dependent FBP biosynthetic enzyme, PfkA, which is a known regulator of Cur and was independently identified in our genetic screen (Figure 1A, B)^8^. We also investigated BT2791 (Upp), a uracil phosphoribosyl transferase putatively involved in nucleotide salvage, because i) the OPPP is required for MdpA-dependent Cur inhibition by generating the essential nucleotide precursors (Figure 4B, C), ii) nutrient limitation is a strong inducer of nucleotide salvage and a condition that activates Cur^19,20^, and iii) Cur contains a predicted nucleotide-binding domain.

### MdpA targets key glycolytic steps to control Cur activity

To explore how MdpA interacts with putative targets, we engineered *wild-type* and Δ*mdpA* strains expressing PyrK or Upp fused to an HA-epitope tag under the control of an aTc-inducible promoter. Strains deficient for Δ*mdpA* produced 2-fold lower HA-tagged PyrK protein amounts compared to *wild-type Bt* (Figure 5E & Supplementary Figure 6A). Similarly, Upp amounts were reduced in Δ*mdpA* compared to *wild-type*, confirming that MdpA regulates the abundance of both enzymes (Figure 5E & Supplementary Figure 6B).

We examined whether PyrK and Upp are required for *mdpA*-dependent Cur inhibition. PyrK is the sole pyruvate kinase in *Bt* unlike *Ec*, which possesses two distinct classes of this enzyme. Thus, genetic examination of a *pyrK* -deficient strain (Δ*pyrK*) was difficult because this mutant did not grow in minimal media containing glucose as a sole carbon source (Supplementary Figure 6C). To circumvent this limitation, we examined Roc amounts in glucose-containing rich media, in which Δ*mdpA* retains elevated Roc production compared to *wild-type* (Figure 5F & Supplementary Figure 6D). Strikingly, Δ*pyrK* exhibited increased Roc that was indistinguishable from Δ*mdpA* or a double mutant (Δ*mdpA* Δ*pyrK*), demonstrating that PyrK and MdpA are epistatic with respect to Roc production and Cur activity (Figure 5F & Supplementary Figure 6D). In contrast, a *upp*-deficient strain (Δ*upp*) did not exhibit increased Roc amounts, indicating that not all MdpA-regulated targets contribute to Cur inhibition, which agrees with the *cur*-independent transcript changes in RNA-seq (Figure 3A & Supplementary Figure 6E, F). Collectively, these data demonstrate that MdpA controls Cur activity in part by regulating the abundance of PyrK, an ATP-producing glycolytic enzyme positioned within the same metabolic pathway as the previously characterized regulator, PfkA (Figure 5G). Combined with the requirement for the OPPP and the diversity of metabolic proteins identified by proximity labeling, these findings indicate that MdpA functions as an important post-translational regulator connecting glucose catabolism to global transcription by controlling Cur.

## Discussion

Organisms across all domains of life have evolved peptidases to regulate glucose homeostasis^1–3^. In this study, we identified MdpA (Figure 1), a conserved M16 class peptidase (Figure 2) that mediates glucose-dependent inhibition of Cur, the master regulator of carbohydrate utilization and an important fitness determinant in the dominant gut genus *Bacteroides* (Figure 3). MdpA specifically facilitates Cur inhibition by glucose, but not fructose (Figure 3), establishing that each monosaccharide engages distinct regulatory pathways, despite being structural isomers and entering glycolysis at adjacent metabolic steps. Importantly, we show that loss of MdpA confers a competitive advantage in gnotobiotic mice when provided a glucose-rich beverage (Figure 3), indicating that this peptidase functions *in vivo* to reduce Cur activity and disrupt *Bacteroides* fitness when glucose is abundant in the host lumen. Additionally, we demonstrate that MdpA controls Cur activity by governing the abundance of the final glycolytic enzyme, PyrK (Figure 5), thereby revealing a glucose-specific regulatory mechanism in *Bacteroides* species that inhibits Cur and silences genes necessary for carbohydrate utilization, intestinal colonization, and host immunotolerance.

MdpA shares conserved motifs with mammalian insulin-degrading enzyme (IDE), which regulates insulin and glucagon levels to balance systemic circulating glucose^1,2^. Our findings further support a hypothesis that M16 peptidases represent a conserved evolutionary approach to glucose homeostasis, despite the phylogenetic distance separating *Bacteroidota, Pseudomonadota*, and *Eukaryota*. Similar to IDE and other M16 peptidases, we show that MdpA can degrade various nonnative substrates, including insulin chain B, when complexed with MdpB, indicating their biochemical activities are shared across kingdoms^27^. Consistent with this notion, our proximity labeling and genetic analyses reveal that MdpA associates with a diverse pool of proteins in glycolysis, nucleotide metabolism, and many other central metabolic pathways. M16 peptidases frequently serve as processing proteases that cleave N-terminal basic amino acid residues from target proteins rather than facilitating their degradation^25^. Indeed, the production of both PyrK and Upp is reduced when *mdpA* is absent, implying that proteolytic activity is important for folding or maturation. This is consistent with the phylogenetic conservation of a putative *cis*/*trans*-prolylisomerase within the same operon as MdpA and its orthologs in *Bacteroides* (Figure 2). Notably, many M16 peptidases, such as *β*-mitochondrial processing peptidases (*β*-MPP)^30^, assume similar roles in other organisms to convert precursors into mature, functional proteins. While our work highlights only a few candidates, MdpA likely facilitates processing of other metabolic enzymes in *Bt* that warrant further examination.

*Bacteroides* species possess ATP- and PPi-dependent glycolytic pathways in place of canonical PTS proteins from Proteobacteria to accomplish sugar catabolism and alter transcription ^7,8^. The identification of *pfkA* in our genetic screen confirms that ATP-dependent glycolysis controls Cur activity in response to simple sugars; however, we now report that glucose and fructose inhibit Cur through separate mechanisms despite converging on the same catabolic pathway. PfkA, which phosphorylates F6P to generate FBP, mediates Cur inhibition by both sugars, likely through FBP production or generating metabolites for cellular energy^8^. In contrast, MdpA is dispensable for Cur inhibition by fructose and acts specifically during glucose catabolism by regulating the abundance of PyrK, the terminal glycolytic enzyme. Because MdpA transcript ^23^ and protein levels do not change depending upon growth condition (Figure 3G & Supplementary Figure 2D), we posit that glucose-dependent regulation occurs at the metabolic level. Examinations of PykA in *Ec*^31^ and *Synechocystis* sp.^32^ have demonstrated G6P and OPPP intermediates, such as ribulose-5P (Ru5P) and ribose-5P (Ri5P) (Figure 4A), as strong allosteric activators of this glycolytic step; however, F6P did not stimulate PykA activity ^31,32^. This provides a potential explanation for glucose-dependent regulation of Cur in *Bacteroides*, whereby glucose, but not fructose, catabolism increases G6P and OPPP intermediates to direct PyrK activity and inhibit Cur. Thus, PfkA and PyrK represent two distinct regulatory nodes in ATP-dependent glycolysis that orchestrate Cur activity when dietary monosaccharides are available.

The evolutionary context of the host diet and microbiome is necessary to understand the consequences of excessive sugar consumption by humans. Gut bacteria have long existed in an intestinal environment where glucose and fructose were transiently encountered during the degradation of dietary and mucosal polysaccharides, rather than as refined monosaccharides. The relatively recent shift toward refined sugars in ultra-processed foods has reshaped the nutrient profile of the gut ecosystem, creating a new evolutionary challenge for bacteria and their hosts^33^. This study deepens our understanding of how a sugar-rich diet has direct consequences for *Bacteroides* by controlling the global regulator, Cur. The deleterious effects of sugar-dependent Cur inhibition are two-fold: first, the *cur*-dependent products Roc and EF-G2 are necessary for bacterial fitness in the mammalian gut^19,21^; second, the host-immunomodulatory signal BT4295 is an important stimulus for T-cell differentiation^17^. Although *Bt* and other *Bacteroides* species exhibit rapid *in vitro* growth on glucose, simple sugar catabolism disrupts colonization and fitness within the context of the human intestine. These findings further reveal a tension between ancestral regulatory logic and a modern host diet, whereby the utilization of glucose for energy reduces fitness determinants necessary for *Bacteroides* colonization and beneficial interactions.

## Materials and Methods

### Bacterial strains and growth conditions

*Ec* strains were cultured aerobically at 37°C on Luria-Bertani (LB) agar or in LB broth. *Bt* strains were cultured at 37°C under anaerobic conditions (85% N_2_, 12.5% CO_2_, 2.5% H_2_) on brain-heart infusion agar (Sigma) containing 5% horse blood (BHI-B). Liquid *Bt* cultures were inoculated into tryptone-yeast extract-glucose (TYG) medium and incubated under anaerobic conditions. Where appropriate, media were supplemented with the following antibiotics: 200 *µ*g/mL gentamicin (Sigma), 2 *µ*g/mL tetracycline (Sigma), or 25 *µ*g/mL erythromycin (Sigma).

### Engineering chromosomal deletions

*Bt* genomic deletions were generated, as previously described^34^, using *pEXCHANGE-tdk* plasmids with flanking sequences that were amplified using primers listed in Supplementary Table 2 and Q5 High-fidelity DNA polymerase (NEB). Resulting amplicons were combined with *pEXCHANGE-tdk* predigested with BamHI-HF (NEB) and SalI-HF (NEB), and assembled using NEBuilder Assembly HiFi (NEB). The resulting constructs were introduced into *Bt* by di-parental mating and chromosomal integration was validated by colony PCR. Parent strains were counter-selected on solid BHI-B medium, unless stated otherwise, containing 200 ng/mL 5-fluoro-2-deoxyuridine (FUdR; DOT Scientific). All strains used in this study were confirmed by Oxford Nanopore of linear amplicons.

### Generation of transposon insertion library

The *Bt* transposon insertion library was generated using pSAM-*Bt*^35^ by di-parental mating. An *Ec* S17-1 donor strain carrying pSAM-*Bt* and the *Bt* recipient strain GT1662 were cultured overnight to stationary phase at 37°C in LB or pre-reduced TYG, respectively. The donor culture was diluted 1:2,000 in LB and the recipient was diluted 1:250 in 10 mL of pre-reduced TYG; both were incubated at 37°C until early exponential phase (OD_600_ = 0.3). Subsequently, 1 mL of donor culture was combined with 10 mL of recipient culture, centrifuged at 7,200× g for 2 minutes, and washed once with 10 mL of 1× phosphatebuffered saline (PBS, Sigma). The pellet was resuspended and plated on BHI-B medium and plates were incubated aerobically at 37°C for 3 hours, followed by overnight incubation under anaerobic conditions to select for *Bt* transconjugants.

The following day, confluent growth was dislodged with a cell spreader into PBS, and the suspension was brought to a final volume of 5 mL in PBS. After homogenization, 1 mL was spread onto BHI-B medium supplemented with 0.2% galactose and appropriate antibiotics. Plates were incubated anaerobically at 37°C for 48 hours. Resulting colonies were pooled into 12 mL PBS containing 20% glycerol, homogenized by vortexing, aliquoted, and stored at −80°C.

### Colony blotting

Aliquots of the *Bt* transposon library were thawed, serially diluted, and plated onto 100 mm Petri dishes containing solid minimal medium supplemented with 0.5% glucose, such that each plate contained approximately 500 colonies. Plates were incubated anaerobically at 37°C for 48 hours, then transferred onto nitrocellulose membranes and subjected to immunoblotting using rabbit anti-HA (Sigma; 1:5,000) and subsequent HRP-conjugated anti-rabbit (GE; 1:5,000), as previously described^20^. Chemiluminescence was detected using ECL Prime Western Blotting Detection Reagent (Cytiva) and imaged on an AI680 (GE). Colonies producing visibly higher chemiluminescent signals were identified, and the corresponding strains were recovered from the master plates, struck for isolation on BHI-B medium, cultured in TYG, and stored at − 80°C. The site of transposon genomic integration was identified using semi-random PCR, as previously described^20^.

### Protein expression and purification

Recombinant proteins were expressed as previously described^8^. Briefly, gene inserts encoding each enzyme were amplified by PCR and introduced into linearized vector using NEBuilder Assembly HiFi (NEB). The resulting plasmids were transformed into chemically competent *Ec* BL21 cells and plated. Overnight cultures grown to stationary phase were diluted 1:50 into fresh LB media. Protein expression was induced by the addition of 500 *µ*M IPTG when cultures reached mid-logarithmic phase (OD_600_ = 0.45–0.5), followed by incubation for 5 hours at 30°C. Cells were harvested by centrifugation and lysed in Lysis Buffer containing 50 mM sodium phosphate (pH 7.4) and 300 mM sodium chloride. Histidine-tagged proteins were purified using Ni^2+^-NTA resin (ThermoFisher) and eluted with Lysis Buffer containing 250 mM imidazole. Proteins were concentrated using centrifugal concentrators (Sigma) and imidazole was removed using Zeba Spin Desalting Columns (ThermoFisher). Additionally, 20% glycerol was added to final protein stocks before storing at − 80°C. Protein concentrations were determined using a BCA assay (ThermoFisher).

### Size-exclusion chromatography

Aliquots of the purified proteins were resolved either individually or after mixing at a 1:1 molar ratio, based on their absorbance at 280 nm, on a Superdex 200 column equilibrated in 10 mM Tris (pH 7.4), 100 mM NaCl, and 0.1 mM TCEP at a flow rate of 0.5 mL/min. The elution profile was monitored in-line at 280 nm. Fractions were collected (0.5 mL) for analysis by SDS-PAGE.

### Western blot

Western blotting was performed as previously described^20^. Stationary phase cultures in TYG were diluted 50-fold into TYG or minimal media containing 0.5% carbon and incubated to midlogarithmic phase (OD_600_ = 0.45–0.65). Cell pellets were resuspended in 400 *µ*L of 1× Tris-buffered saline (TBS) containing 1 mM EDTA (Sigma) and 0.5 mg/mL chicken egg white lysozyme (Sigma) and lysed using a Mini-Beadbeater (BioSpec). Lysates were clarified by centrifugation and total protein concentration in the supernatant was determined by measuring absorbance at 280 nm using a NanoQuant plate in a Spark microplate reader (Tecan). Samples corresponding to 30 *µ*g of total protein were combined with 4 *µ*L of 4× LDS sample buffer (ThermoFisher) containing 100 mM dithiothreitol (DTT), boiled, and resolved on 4–12% Bis-Tris NuPAGE gel (ThermoFisher) in MES running buffer (Invitrogen). Proteins were transferred to nitrocellulose membranes using an iBlot2 transfer apparatus (Invitrogen). Following transfer, membranes were blocked for 1 hour at room temperature in TBS containing 3% skim milk (BD), then probed with either rabbit anti-HA (Sigma; 1:5,000) or rabbit anti-GroEL (Sigma; 1:5,000), followed by HRP-conjugated anti-rabbit (Cytiva; 1:5,000). Chemiluminescence was detected using ECL Prime Western Blotting Detection Reagent (Cytiva) and imaged on an AI680 (GE). Band intensities were quantified using Image Studio Lite software (LI-COR Biosciences).

### Phylogenetic tree construction

Protein sequences were retrieved from the Orthologous Matrix (OMA) database for two orthologous groups: 946369 and 946368. For each group, member sequence and genomic metadata were obtained using the OmaDB R package. Protein sequences were aligned using MUSCLE via the Bioconductor msa package (v. 1.40.0). Maximum-likelihood tree inference was performed with IQ-TREE 2 (v. 2.3.6) using ModelFinder for automatic model selection (LG+R8) and the default tree search strategy. The resulting tree was midpoint-rooted for visualization using ggtree and Circos.

### Enzyme assays

Proteolytic assays were performed as previously described^24^ with the following modifications. All assays were conducted in 50 mM Tris (pH 7.4) containing 3 mM MgCl_2_. All assays were performed at final concentration of 0.1 mg/mL of enzyme. Cleavage of fluorogenic peptide Abz-pAK (GenScript) was measured kinetically for 10 minutes, and activity was quantified using the change in slope compared to substrate alone. Proteolysis of insulin chain B and dynorphin A were analyzed using a C-18 Agilent Poroshell column fitted to an Agilent 1260 Infinity II. Peptides were eluted using a 17-minute linear gradient from 10% acetonitrile:90% H_2_O to 90% acetonitrile:10% H_2_O, with both eluants containing 0.1% formic acid. Chromatograms were analyzed using OpenLab Chemstation software (Agilent).

### qPCR

Stationary-phase *Bt* cultures were diluted 1:50 into pre-reduced minimal media containing 0.5% carbon and incubated to midlogarithmic phase (OD_600_ = 0.45–0.60). 1 mL of culture was pelleted by centrifugation, flash-frozen on dry ice, and stored at −80°C. RNA was extracted using the RNeasy Mini Kit (QIAGEN), quantified by spectrometry, and 1 *µ*g was reverse transcribed into cDNA using the SuperScript IV VILO Master Mix with ezDNase (Invitrogen), according to the manufacturer’s directions. Transcript levels were quantified using a QuantStudio 5 Real-Time PCR system (ThermoFisher) with PowerUp SYBR Green Master Mix (ThermoFisher) and gene-specific primers (Supplementary Table 2). The 16S rRNA gene (*rrs*) was used as the internal reference for normalization in all qPCR analyses.

### RNA sequencing

RNA-seq was performed as previously described^19^ with the following modifications. *Bt* cultures were grown to mid-logarithmic phase (OD_600_ = 0.45–0.60) in minimal medium containing 0.5% carbon as described for qRT-PCR. 10 mL of each culture was harvested by centrifugation and stabilized using 10 mL of RNAprotect Bacterial Reagent (Qiagen; diluted 2:1 with nuclease-free water), then frozen on dry ice and stored at − 80°C. Total RNA was isolated using the RNeasy Mini Kit (Qiagen) with on-column DNase I treatment. Eluted RNA was treated with Turbo DNase (Invitrogen) for 30 min at 37°C to remove residual genomic DNA and purified again using the RNeasy Mini Kit. Strand-specific cDNA libraries were prepared by Azenta Life Sciences following assessment of RNA integrity using a Bioanalyzer (Agilent). Libraries were sequenced on an Illumina HiSeq 4000 platform, generating approximately 50 million 150-bp paired-end reads per sample. Sequencing reads were aligned to the *Bt* genome (GenBank accession NC_004663) using Bowtie2 (v2.3.4) on the Galaxy web server. Read quantification was performed with featureCounts (v1.6.3), and differential gene expression analysis was carried out using DESeq2 (v1.18.1) with an adjusted *P*-value threshold of 0.05. Gene annotations and expression values for all transcripts are compiled in Supplementary Table 4.

### BioID sample preparation and streptavidin affinity purification

Stationary-phase *Bt* cultures harboring aTc-inducible MdpA-BirA, E50Q-BirA, or BirA-alone constructs were diluted 1:50 into pre-reduced minimal medium containing 0.5% glucose and incubated anaerobically at 37°C. At early mid-logarithmic phase (OD_600_ = 0.45–0.60), anhydrotetracycline (aTc) was added to a final concentration of 100 *µ*g/mL. Cell pellets were resuspended in 1 mL of freshly prepared lysis buffer (8 M urea, 50 mM Tris (pH 7.4), 1 mM DTT, and Halt Protease Inhibitor Cocktail, EDTA-free (ThermoFisher)) and lysed using a Mini-Beadbeater (BioSpec).

For streptavidin affinity purification, 150 *µ*L of Streptavidin Sepharose High Performance beads (Cytiva) were equilibrated by washing three times with 1 mL of lysis buffer. Filtered lysates were added to the equilibrated resin and incubated overnight at 4°C with gentle rocking. Resin-bound samples were pelleted by centrifugation and the supernatant was removed. Beads were washed three times with 1 mL of wash buffer (8 M urea, 50 mM Tris, pH 7.4) for 10 minutes at room temperature. An aliquot of the washed resin was reserved for quality control, and the remaining beads were stored at −80°C until proteomic analysis.

### Proteomic and pathway enrichment analysis

Protein mass spectrometry was performed by Creative Proteomics. Isolated protein samples from BirA, MdpA-BirA, or E50Q-BirA were matched to the Bacteroides thetaiotaomicron VPI-5482 proteome. Relative abundance of proteins from each fusion construct were normalized to samples from the BirA control. Abundance score represents the mean relative protein amounts between both fusion constructs. KEGG pathway analysis was performed in R using the *Bt* database (bth). Proteins identified in the proteomics dataset were trimmed to have a minimum gene ratio threshold of 0.5% and gene count of 2, then matched against pathways excluding global and overview maps (IDs: 01100, 01110, 01120, 01200, 01210, 01212, 01230, 01232, 01250, 01240, and 01320). The gene ratio was calculated as the number of matched proteins divided by the total number of proteins in each pathway. The resulting dot-plots were visualized using the ggplot2 R package.

## Acknowledgements

We thank Donalee McElrath, Michelle Irish, Jacob Perryman, and Alexandra Sprinkle for assistance with germ-free mouse experiments. Additionally, we thank Jennifer Modesto, Jennifer Lausch, Jessica Gaydos, and Aayush Gupta for helpful comments during manuscript preparation. This work was supported, in part, by National Institutes of Health Grants DK132711 and GM147178 to G.E.T. and GM123798 to E.A.G.

## Supplementary Information

**Supplementary Figure 1.**
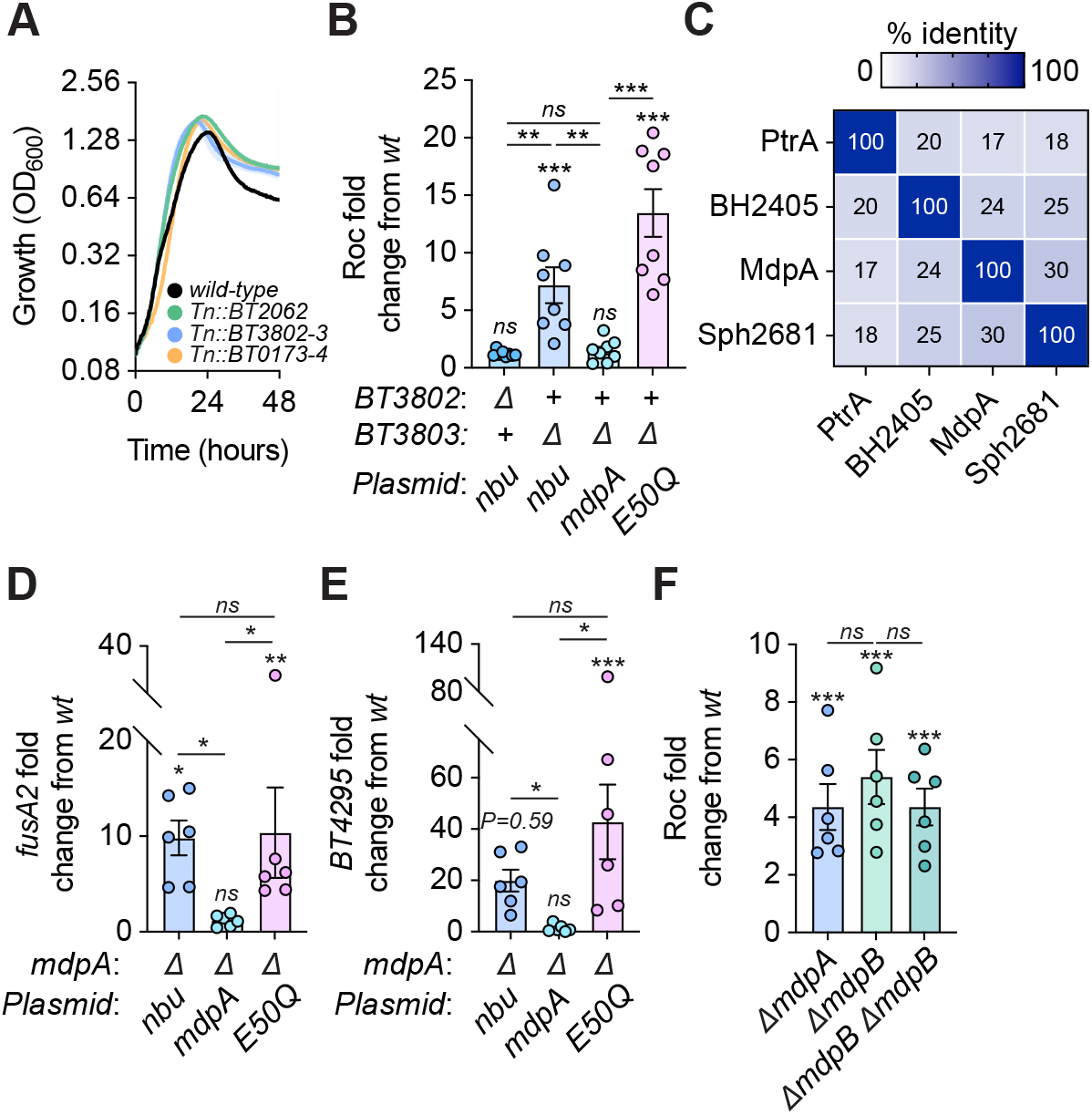
MdpA requires a conserved glutamate residue for glucose-dependent Roc silencing. **A** Growth of *wild-type, BT2062*::Tn, *BT3802-3*::Tn, or *BT0173-4*::Tn in minimal media containing glucose. **B** Fold change of Roc protein amounts relative to *wild-type* from Δ*BT3802* or Δ*mdpA* harboring an empty vector (nbu) or a complementing plasmid grown in minimal media containing glucose. **C** Percent identity matrix of MdpA with M16 peptidases in related bacterial species. **D, E** Fold change of *fusA2* (**D**) or *BT4295* (**E**) transcript amounts relative to *wild-type* from Δ*mdpA*, Δ*cur*, or Δ*mdpA* Δ*cur* complemented with an empty vector (nbu), *mdpA*, or E50Q mutant grown in minimal media containing glucose. **F** Fold change of Roc protein amounts relative to *wild-type* from Δ*mdpA*, Δ*mdpB*, or Δ*mdpA* Δ*mdpB* grown in minimal media containing glucose. For **A**, *n* = 8 biological replicates; error is SEM in color matched shading. For **B**, *n* = 8 biological replicates; error is SEM. For **D, E, F**, *n* = 6 biological replicates; error bars is SEM. *P*-values were calculated using one-way ANOVA and * represent values *<* 0.05, ** *<* 0.01, *** *<* 0.001.

**Supplementary Figure 2.**
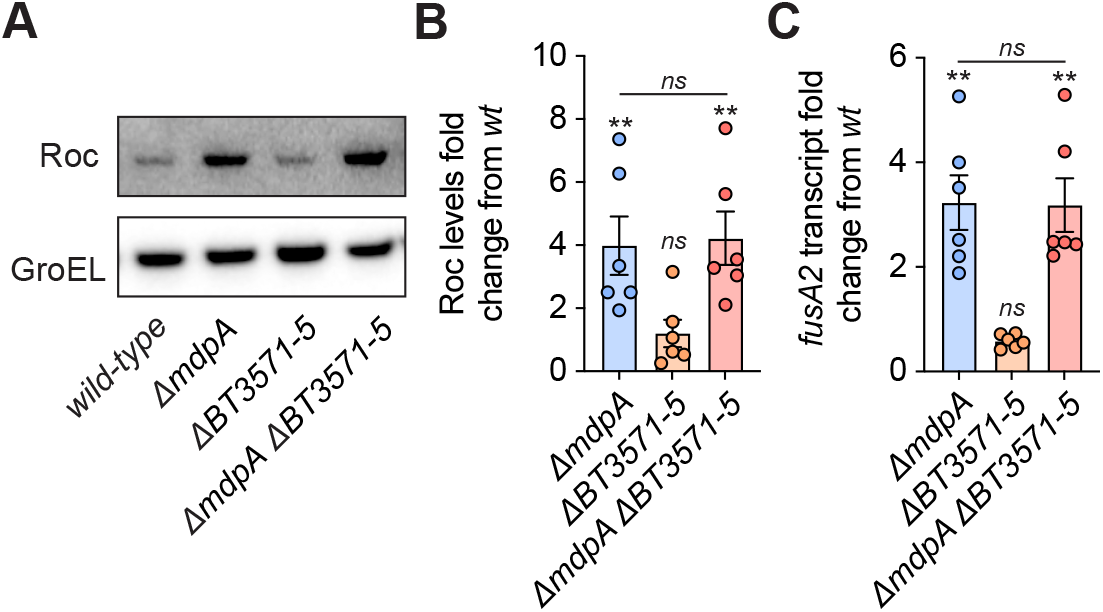
The *BT3571-5* locus does not control Cur activity. **A** Western blot of Roc protein amounts from *wild-type*, Δ*mdpA*, Δ*BT3571-5*, or Δ*mdpA* Δ*BT3571-5* grown in minimal media containing glucose. **B** Fold change of Roc protein amounts relative to *wild-type* from Δ*mdpA*, Δ*BT3571-5*, or Δ*mdpA* Δ*BT3571-5* grown in minimal media containing glucose. **C** Fold change of *fusA2* transcript amounts relative to *wild-type* from Δ*mdpA*, Δ*BT3571-5*, or Δ*mdpA* Δ*BT3571-5* grown in minimal media containing glucose. For **B, C**, *n* = 6 biological replicates; error is SEM. *P*-values were calculated using one-way ANOVA and ** represents values *<* 0.01.

**Supplementary Figure 3.**
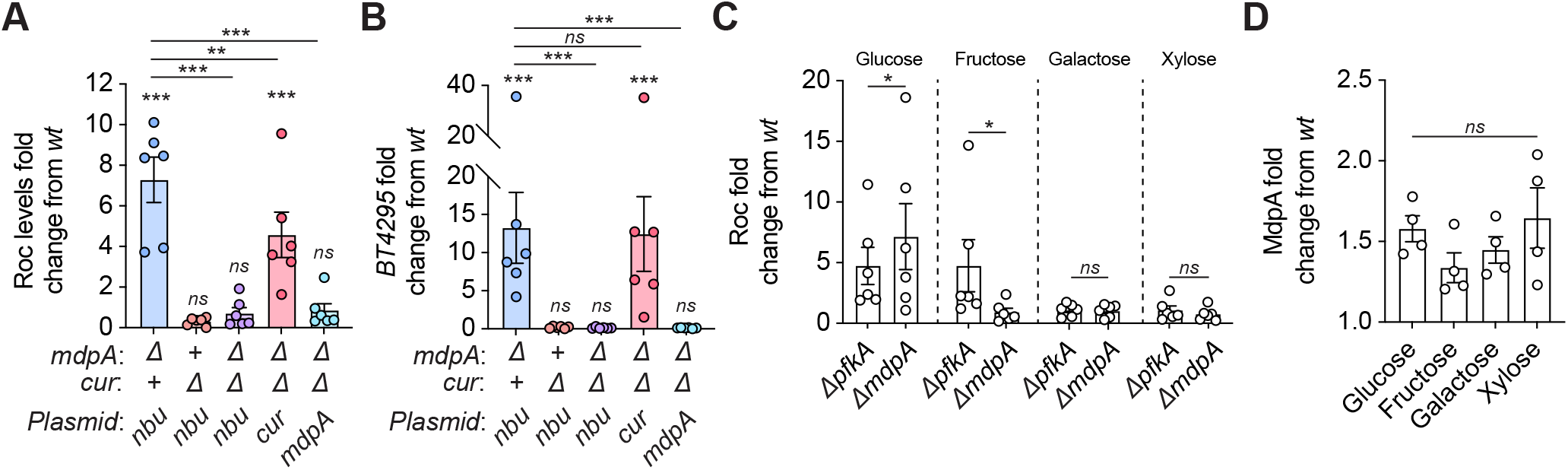
*mdpA* is required for Roc silencing in glucose but not fructose. **A** Fold change of Roc protein amounts relative to *wild-type* from Δ*mdpA*, Δ*cur*, or Δ*mdpA* Δ*cur* harboring an empty vector (nbu) or complementing plasmids grown in minimal media containing glucose. **B** Fold change of *BT4295* transcript amounts relative to *wild-type* from Δ*mdpA*, Δ*cur*, or Δ*mdpA* Δ*cur* harboring an empty vector (nbu) or complementing plasmids grown in minimal media containing glucose. **C** Fold change of Roc protein amounts relative to *wild-type* from Δ*pfkA* or Δ*mdpA* mutants grown in minimal media containing glucose, fructose, galactose, or xylose. **D** Fold change of MdpA protein amounts relative to *wild-type* from Δ*mdpA* grown in minimal media containing glucose, fructose, galactose, or xylose. For **A, B, C**, *n* = 6 biological replicates; error is SEM. For **D**, *n* = 4 biological replicates; error is SEM. *P*-values were calculated using one-way ANOVA and * represents values *<* 0.05, ** *<* 0.01, and *** represents values *<* 0.001.

**Supplementary Figure 4.**
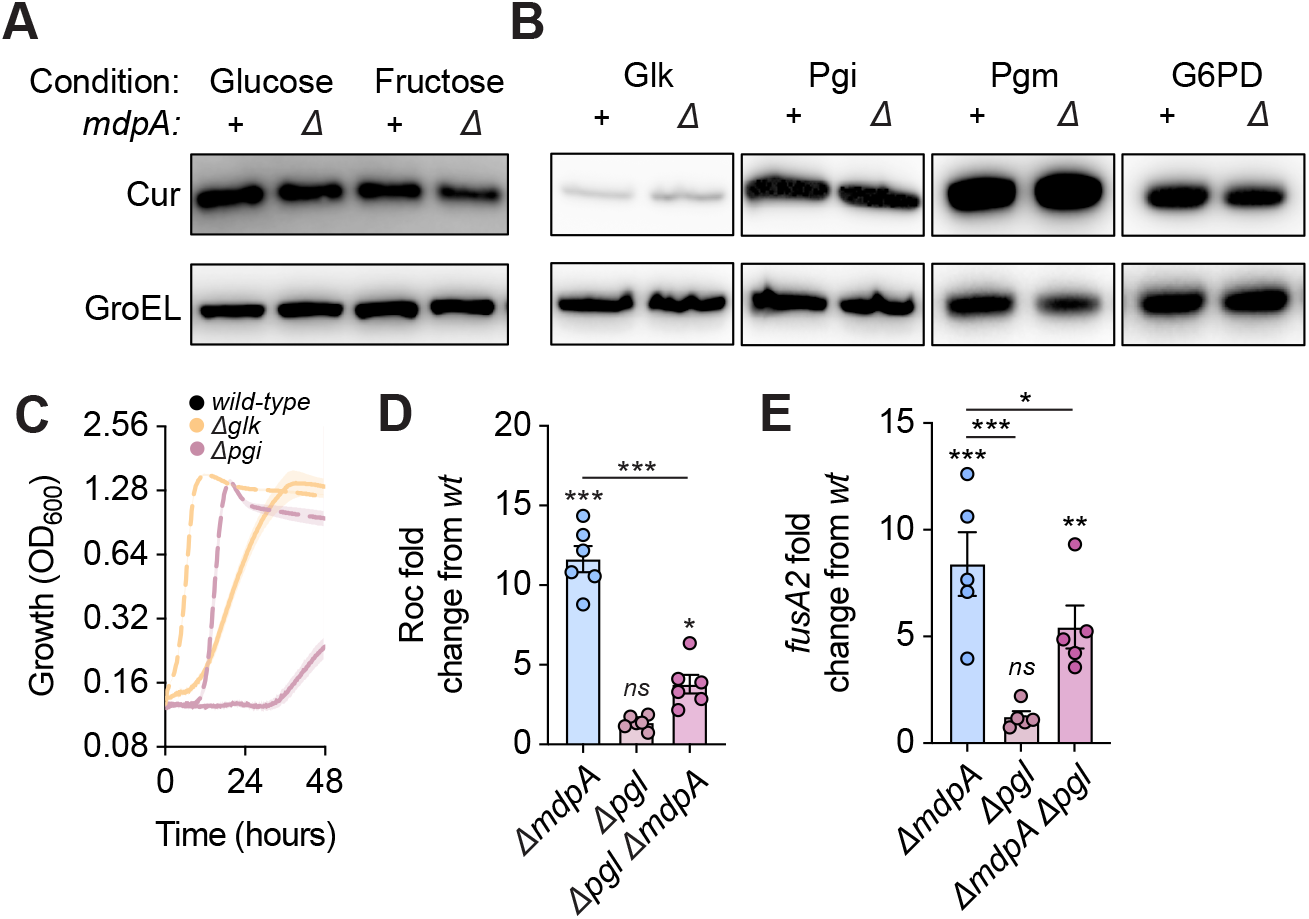
The oxidative pentose phosphate pathway is required for MdpA control of Cur activity. **A** Western blot of Cur protein amounts from *wild-type* or Δ*mdpA* grown in minimal media containing glucose (left) or fructose (right). **B** Western blot of Glk, Pgi, Pgm, and G6PD protein amounts from *wild-type* or Δ*mdpA* grown in minimal media containing glucose. **C** Growth of *wild-type*, Δ*glk*, or Δ*pgi* strains in rich media containing glucose (solid line) or fructose (dashed line). **D** Fold change of Roc protein amounts relative to *wild-type* from Δ*mdpA*, Δ*pgl*, or Δ*mdpA* Δ*pgl* grown in minimal media containing glucose. **E** Fold change of *fusA2* transcript amounts relative to *wild-type* from Δ*mdpA*, Δ*pgl*, or Δ*mdpA* Δ*pgl* grown in minimal media containing glucose. For **C**, *n* = 4 biological replicates; error is SEM in color matched shading. For **D**, *n* = 4 biological replicates, error is SEM. For **E**, *n* = 6 biological replicates, error is SEM. For **D, E**, *P*-values were calculated using one-way ANOVA and * represents values *<* 0.05, ** *<* 0.01, *** *<* 0.001.

**Supplementary Figure 5.**
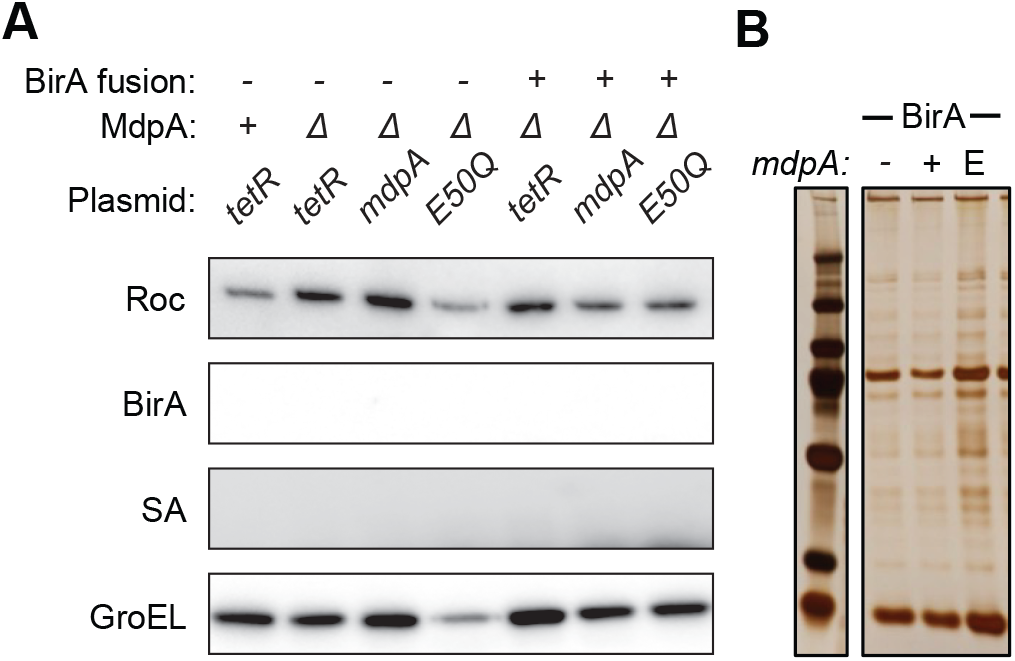
Identification of MdpA targets using biotinylation. **A** Western blot of Roc, BirA, and SA protein amounts in the absence of aTc-inducible expression of MdpA or E50Q-BirA fusion proteins from *wild-type* or Δ*mdpA* grown in minimal media containing glucose. **B** Silver staining of elutions following aTc-induction and streptavidin affinity purification from strains harboring an empty vector, *mdpA*, or *E50Q* grown in minimal media containing glucose.

**Supplementary Figure 6.**
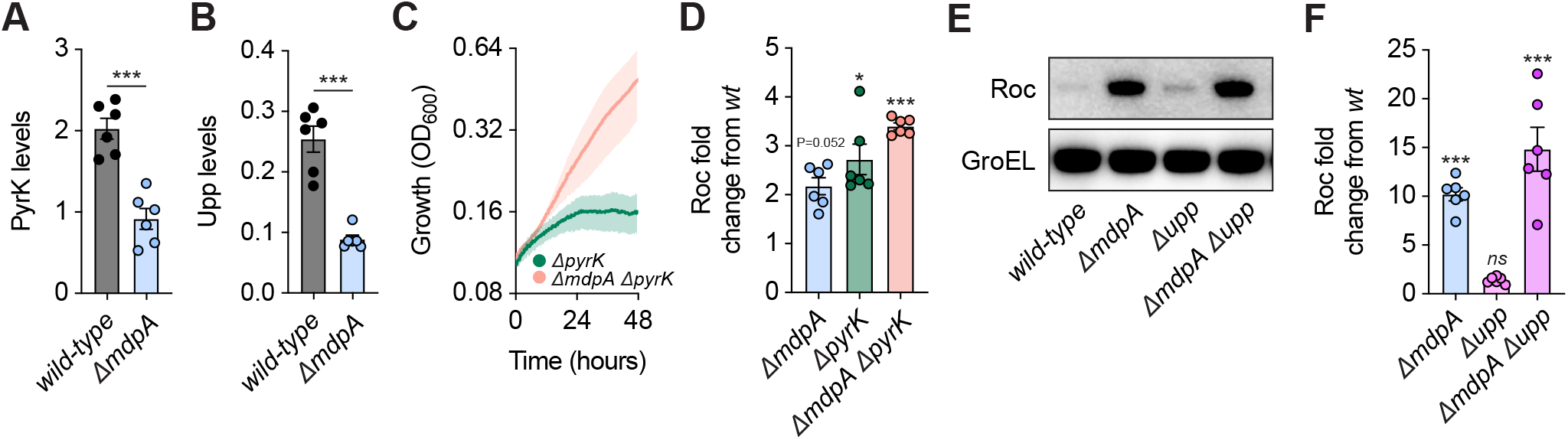
MdpA governs the abundance of proteins that mediate glucose-dependent Cur inhibition. **A, B** Quantification of PyrK (**A**) or Upp (**B**) protein amounts in *wild-type* or Δ*mdpA* grown in minimal media containing glucose. **C** Growth of Δ*pyrK* or Δ*mdpA* Δ*pyrK* in minimal media containing glucose. **D** Fold change of Roc protein amounts relative to *wild-type* from Δ*mdpA*, Δ*pyrK*, or Δ*mdpA* Δ*pyrK* grown in minimal media containing glucose. **E** Western blot of Roc protein amounts from *wild-type*, Δ*mdpA*, Δ*upp*, or Δ*mdpA* Δ*upp* grown in minimal media containing glucose. **F** Fold change of Roc protein amounts relative to *wild-type* from Δ*mdpA*, Δ*upp*, or Δ*mdpA* Δ*upp* grown in minimal media containing glucose. For **A, B, D, F**, *n* = 6 biological replicates; error is SEM. For **C**, *n* = 8 biological replicates; error is SEM in color matched shading. For **A** and **B**, *P*-values were calculated using an unpaired t-test and *** represents values *<* 0.001. For **D** and **F**, *P*-values were calculated using one-way ANOVA and * represents values *<* 0.05, ** *<* 0.01, *** *<* 0.001.

